# Protective efficacy of rhesus adenovirus COVID-19 vaccines against mouse-adapted SARS-CoV-2

**DOI:** 10.1101/2021.06.14.448461

**Authors:** Lisa H. Tostanoski, Lisa E. Gralinski, David R. Martinez, Alexandra Schaefer, Shant H. Mahrokhian, Zhenfeng Li, Felix Nampanya, Huahua Wan, Jingyou Yu, Aiquan Chang, Jinyan Liu, Katherine McMahan, Kenneth H. Dinnon, Sarah R. Leist, Ralph S. Baric, Dan H. Barouch

## Abstract

The global COVID-19 pandemic has sparked intense interest in the rapid development of vaccines as well as animal models to evaluate vaccine candidates and to define immune correlates of protection. We recently reported a mouse-adapted SARS-CoV-2 virus strain (MA10) with the potential to infect wild-type laboratory mice, driving high levels of viral replication in respiratory tract tissues as well as severe clinical and respiratory symptoms, aspects of COVID-19 disease in humans that are important to capture in model systems. We evaluated the immunogenicity and protective efficacy of novel rhesus adenovirus serotype 52 (RhAd52) vaccines against MA10 challenge in mice. Baseline seroprevalence is lower for rhesus adenovirus vectors than for human or chimpanzee adenovirus vectors, making these vectors attractive candidates for vaccine development. We observed that RhAd52 vaccines elicited robust binding and neutralizing antibody titers, which inversely correlated with viral replication after challenge. These data support the development of RhAd52 vaccines and the use of the MA10 challenge virus to screen novel vaccine candidates and to study the immunologic mechanisms that underscore protection from SARS-CoV-2 challenge in wild-type mice.

**Importance:** We have developed a series of SARS-CoV-2 vaccines using rhesus adenovirus serotype 52 (RhAd52) vectors, which exhibits a lower seroprevalence than human and chimpanzee vectors, supporting their development as novel vaccine vectors or as an alternative Ad vector for boosting. We sought to test these vaccines using a recently reported mouse-adapted SARS-CoV-2 (MA10) virus to i) evaluate the protective efficacy of RhAd52 vaccines and ii) further characterize this mouse-adapted challenge model and probe immune correlates of protection. We demonstrate RhAd52 vaccines elicit robust SARS-CoV-2-specific antibody responses and protect against clinical disease and viral replication in the lungs. Further, binding and neutralizing antibody titers correlated with protective efficacy. These data validate the MA10 mouse model as a useful tool to screen and study novel vaccine candidates, as well as the development of RhAd52 vaccines for COVID-19.

## Introduction

A critical component of the evaluation of vaccine candidates for COVID-19 has been the development of pre-clinical challenge models. Transgenic mice [1-5], hamsters [6-9], and non-human primates [10-12] have been shown to support viral replication and, to varying degrees, clinical disease following infection with SARS-CoV-2 [13]. We recently described a mouse-adapted virus (MA10) to enable challenge of standard, wild-type laboratory mice and recapitulate several key features of human disease, such as viral replication in respiratory tract tissues and severe infection-associated weight loss [14]. This model has been explored to evaluate small molecule antivirals and candidate monoclonal antibodies for prophylactic or therapeutic applications, as well as prototype vaccine candidates [14-17]. For example, initial studies using viral replicon particles expressing SARS-CoV-2 Spike protein demonstrate the capacity of vaccines to restrain MA10 infection and disease [14].

This model has not yet been utilized to study the characteristics of vaccine-elicited immune responses that protect against clinical disease and viral replication. Thus, we sought to test a series of candidate rhesus adenovirus serotype 52 (RhAd52) [18] vector-based vaccines expressing engineered versions of SARS-CoV-2 Spike. RhAd52 vectors have lower seroprevalence in human populations than Ad26 vectors, which recently received FDA Emergency Use Authorization as a COVID-19 vaccine [19, 20]. We aimed to probe correlates of protection, including whether similar immune parameters such as neutralizing antibody titers emerge in the mouse model as predictors of challenge outcome, as has been observed in hamsters and nonhuman primates [9, 10, 21]. These data will inform applications of the MA10 virus to study key questions about clinical disease, infection, or both.

## Results

### Immunogenicity of RhAd52 vectors

We designed a series of replication incompetent viral vector vaccines using rhesus adenovirus serotype 52 (RhAd52) vectors [18, 22] that encode variations of the SARS-CoV-2 Spike (S) protein (Fig 1A). Similar to our previous reports with human adenovirus serotype 26 (Ad26) vectors [9, 23, 24], inserts included: i) unmodified S, ii) truncations of the cytoplasmic tail (S.dCT) or the transmembrane region (S.dTM), or iii) select fragments, including the S1 domain and the receptor binding domain (RBD). In some cases, immunogens were modified with mutation of the furin cleavage site and the addition of proline mutations to stabilize protein prefusion conformation (PP) [25-27]. To explore the potential of candidate RhAd52 vaccines to elicit humoral immune responses, groups of wild-type BALB/c mice were immunized with 10^9^ viral particles (VPs) of these vaccines via the intramuscular route (Fig 1B). Peripheral blood was collected on a biweekly basis to monitor antibody responses in serum.

**Fig 1.**
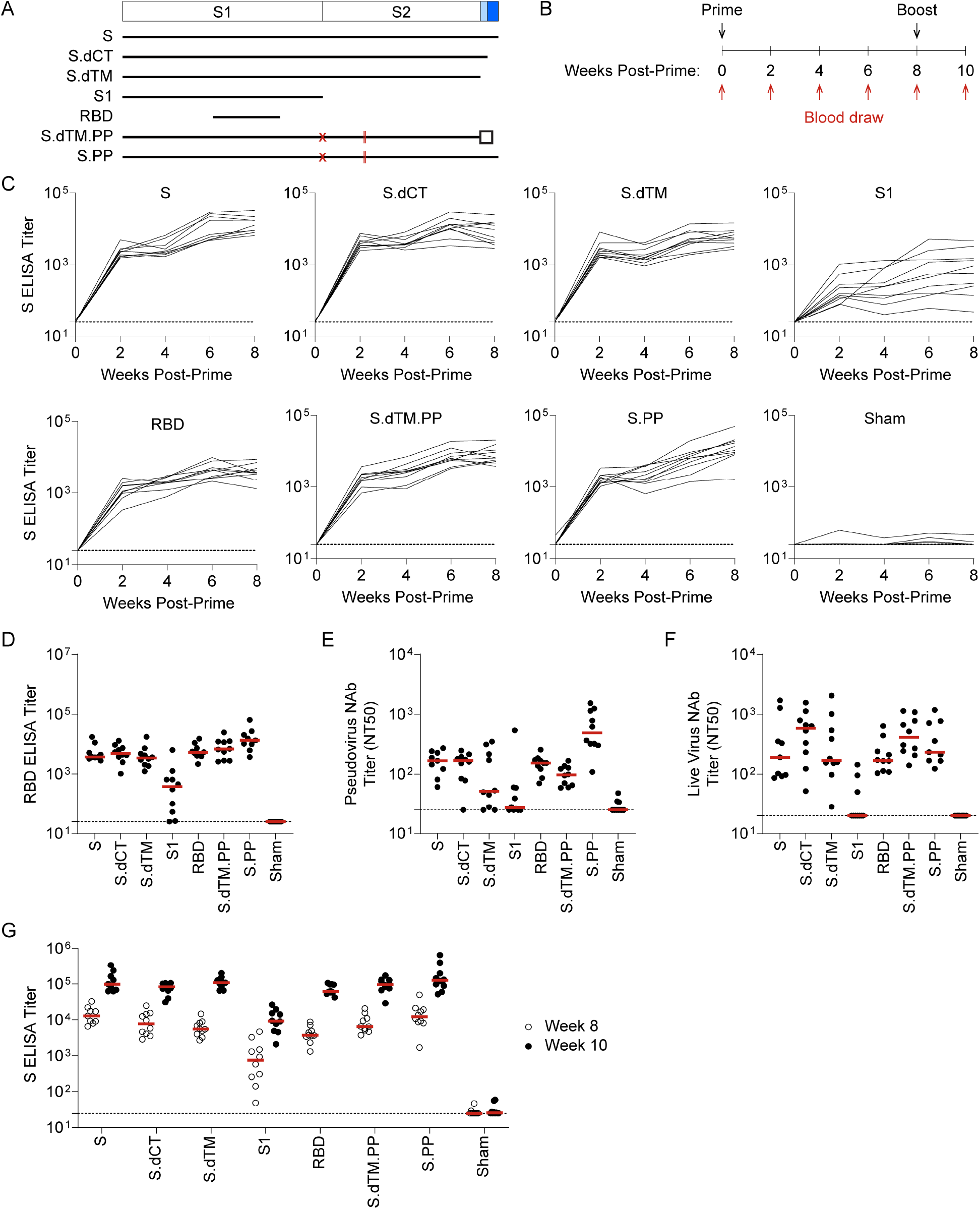
RhAd52 vaccines elicit robust S-specific binding antibody responses. A) A series of replication incompetent RhAd52 vectors, encoding for variations on the SARS-CoV-2 Spike (S) protein, was designed. Inserts included i) S, ii) deletion of the cytoplasmic tail (S.dCT), iii) deletion of the transmembrane domain and cytoplasmic tail (S.dTM), iv) the S1 domain with a foldon trimerization tag, v) the receptor binding domain (RBD) a foldon trimerization tag, vi) deletion of the transmembrane domain and cytoplasmic tail, with mutation of the furin cleavage site (red X), addition of stabilizing proline mutations (red lines), and a foldon trimerization tag (S.dTM.PP), and vii) S with mutation of the furin cleavage site (red X) and addition of stabilizing proline mutations (red lines) (S.PP). B) To explore the immunogenicity of these vaccine candidates, wild-type BALB/c mice were immunized at week 0 with 10^9^ viral particles (VPs) of candidate RhAd52 vaccines or sham. Peripheral blood was collected at baseline and every two weeks following vaccination to monitor antibody responses in serum. Eight weeks post-prime, mice were administered a homologous boost to explore the potential to boost responses. C) For each RhAd52 insert, as well as sham controls, S-specific binding antibody responses were quantified through enzyme-linked immunosorbent assay (ELISA) in serum every two weeks post-prime. D) The distribution of RBD-specific ELISA titers at week 8 across candidate RhAd52 vectors. Red lines indicate the median titer of each group. Neutralizing activity of vaccine-elicited antibody responses were assessed through E) pseudovirus or F) live SARS-CoV-2 virus *in vitro* neutralization assays. The 50% neutralization titer (NT50) is displayed, with the median of each vaccine regimen indicated with a red line. G) The distribution of S-specific ELISA titers at week 8 (open circles) and week 10 (two weeks post-boost, closed circles) was measured to characterize the immunogenicity of a homologous boost with candidate RhAd52 vaccines. Red lines indicate the median titer of each group. For panels C-G, N=9-10 mice/group and endpoint binding titers are reported. Representative data from one of two similar experiments are shown.

At week 2 following the initial immunization, 100% of mice immunized with RhAd52 vaccines, irrespective of the immunogen insert, exhibited SARS-CoV-2 S-specific binding antibodies by enzyme-linked immunosorbent assay (ELISA) (Fig 1C). These responses generally increased over the time frame of 2-8 weeks post-prime. At week 8, RBD-specific binding responses were also observed in 100% of mice immunized with RhAd52 candidates (Fig 1D). Furthermore, antibody function was assessed using *in vitro* assays to quantify the potential to neutralize either a pseudotyped virus [10, 28, 29] (Fig 1E) or live SARS-CoV-2 virus [10, 28, 30, 31] (Fig 1F). Similar to the binding results, neutralizing titers were elicited by several of the RhAd52 candidate vaccines, with the lowest responses observed following immunization with the S1 domain insert, in which a subset of mice exhibited no detectable neutralizing responses at week 8. Mice received a second identical dose of the respective RhAd52 vectors at week 8. Two weeks following the boost immunization, median S-specific ELISA titers were found to increase by approximately 10-fold for all the vaccine candidates (Fig 1G). These data demonstrate the immunogenicity of a homologous boost with a second immunization of a RhAd52 vector.

We next designed a series of immunization regimens that we hypothesized would i) allow direct comparison of protective efficacy of single-shot versus two-dose prime-boost schedules, and ii) generate a range of binding and neutralizing antibody responses that could enable analyses of correlates of protection following challenge [21, 24, 28]. Briefly, groups of mice were immunized with a prime and a matched boost with the seven candidate RhAd52 vaccines, as in Fig 1B. At the time of boost (i.e., week 8), additional groups of mice were immunized with a single dose of select vaccines, RhAd52.S, RhAd52.S.dCT, and RhAd52.S.PP. At week 12, serum was collected to assess antibody responses prior to viral challenge. Expansion of S-specific (Fig 2A) and RBD-specific (Fig 2B) binding antibody titers was again observed in all vaccinated mice. Groups of mice that received the two-dose regimens exhibited approximately one-log higher median titers compared with groups administered a single immunization. Furthermore, consistent with our previous data using a DNA vaccination platform in non-human primates [28], the S1 insert drove the lowest binding responses.

**Fig 2.**
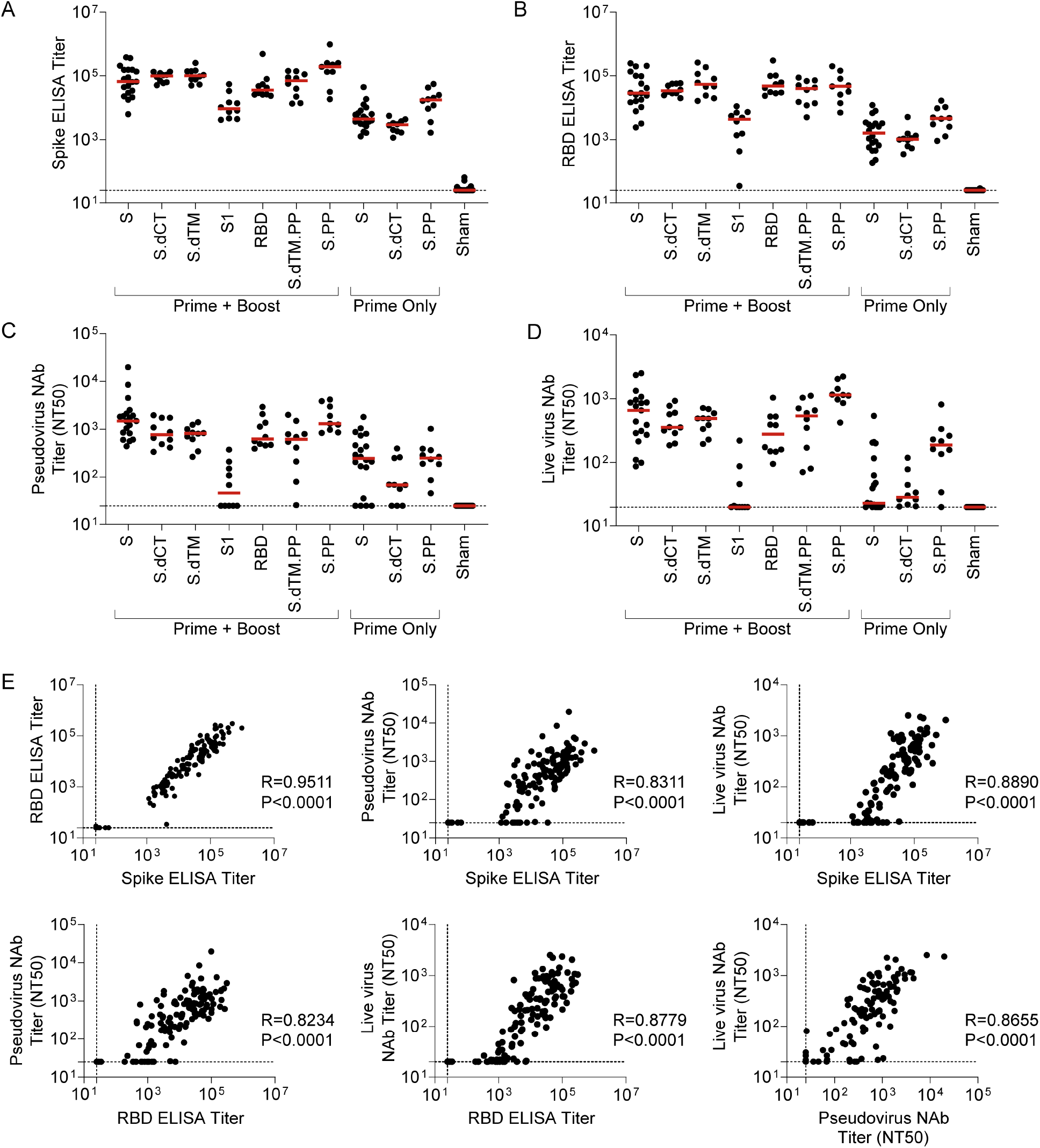
Serum binding and neutralizing antibody responses are tightly linked following RhAd52 vaccination. Groups of mice were administered a prime (week 0) and a boost (week 8) of 10^9^ VP the indicated RhAd52 vaccines. At the time of boost (i.e., week 8), additional groups of mice were administered a single dose (i.e., Prime Only) of 10^9^ VP of select indicated RhAd52 vaccines. At week 12, serum was analyzed. A) S-specific and B) RBD-specific ELISA titers are shown, with median titer for each regimen indicated with a red line. Neutralizing activity of vaccine-elicited antibody responses were assessed through C) pseudovirus or D) live SARS-CoV-2 virus neutralization assays. The 50% neutralization titer (NT50) is displayed, with the median of each vaccine regimen indicated with a red line. E) Spearman correlation analyses of binding and neutralizing antibody responses are displayed. For panels A-E, data are pooled from two similar experiments. For panels A-D, N=9-10 mice/group for all regimens, with the exception of RhAd52.S Prime + Boost (N=19), RhAd52.S Prime Only (N=20), and Sham (N=23). In Panel E, data from all vaccine regimens are pooled to explore the relationship between binding and neutralizing antibody function independent of the RhAd52 insert.

In evaluating neutralizing antibody activity elicited by these vaccine regimens, similar patterns were observed with pseudovirus (Fig 2C) and live virus (Fig 2D) assays. Among the groups administered prime-boost dosing schedules, high neutralizing antibody titers were observed across all vaccines with the exception of the S1 immunogen, for which 50% neutralization titer (NT50) values were below the assay limit of detection for several mice. Groups that received only one dose of select inserts exhibited lower neutralizing titers compared with boosted mice, with detectable responses in a subset of mice that, on average, were approximately one log lower than the boosted groups that received full length or truncated (e.g., S.dCT) vaccines. As consistent trends were observed across binding and neutralizing titers in the relative magnitude of responses elicited by various candidate vaccines, correlation analyses were performed to assess the relationship between these immunologic readouts (Fig 2E). Highly significant (P<0.0001, Spearman correlation) strong positive correlations were observed between binding ELISA titers to S and RBD proteins and capacity to neutralize either pseudovirus or live SARS-CoV-2 virus.

### Protective efficacy of RhAd52 vector vaccines against MA10 challenge

At week 12, all groups of mice were challenged to evaluate whether vaccine-elicited responses protected from clinical disease and viral replication in this mouse-adapted model [14]. Vaccinated mice were challenged on day 0 with 10^4^ PFU SARS-CoV-2 MA10 via the intranasal route (Fig 3A). Half of the mice were followed through day 4 post-challenge and body weight was monitored daily for signs of clinical disease. At the terminal time point, lung tissue was collected, and outgrowth assays were performed to quantify replication-competent virus (i.e., plaque forming units (PFU) in this key respiratory tract tissue. In parallel, half of the mice were sacrificed at day 2 post-challenge to measure virus in the lungs. We hypothesized this approach would allow evaluation of the potential to restrain clinical symptoms of disease as well as enable a virologic endpoint at the time of typical peak viral load (i.e., day 2 post-challenge).

**Fig 3.**
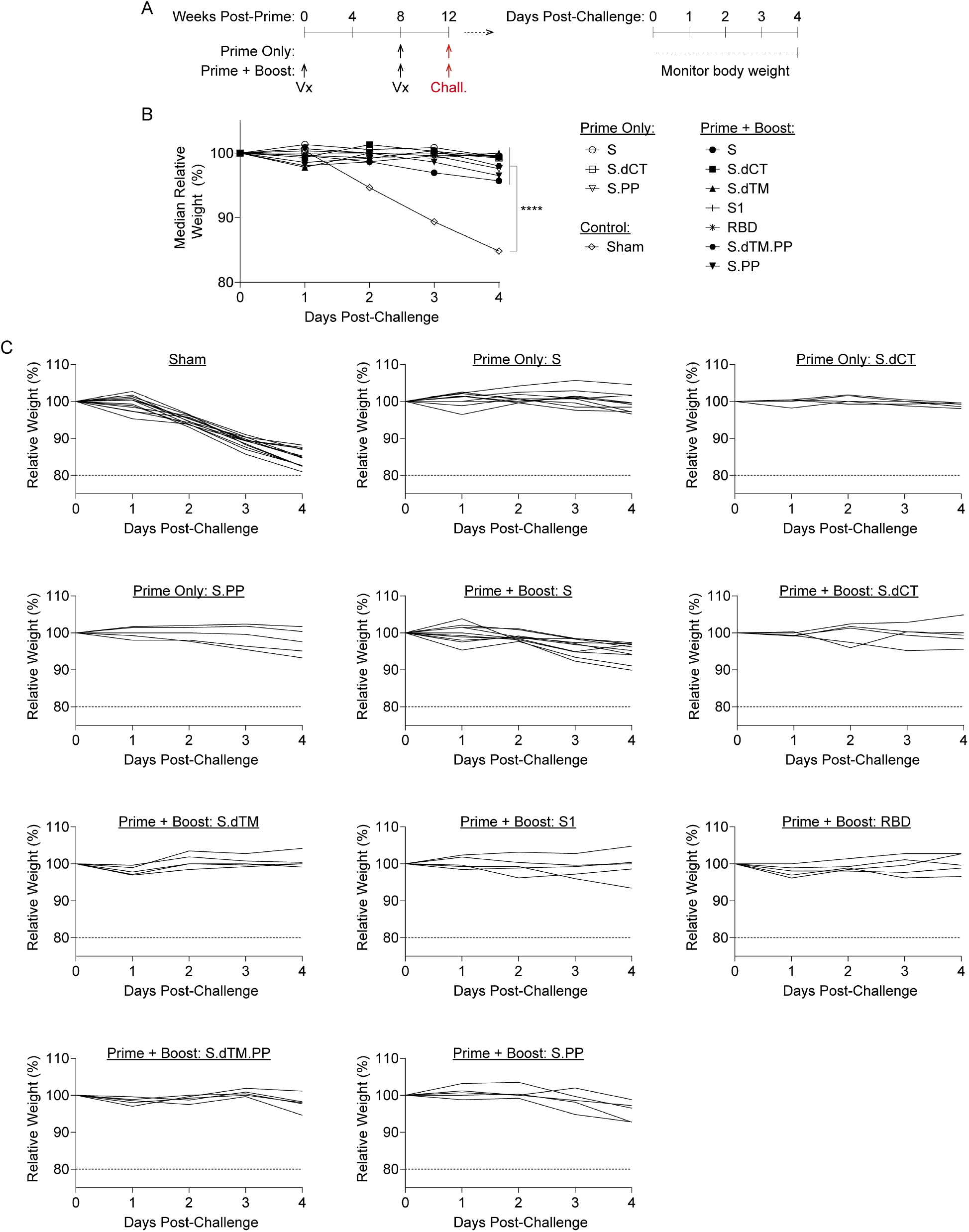
RhAd52 vaccines protect from clinical disease following mouse-adapted SARS-CoV-2 challenge. A) Groups of mice were immunized with either a prime and boost or a prime only of 10^9^ viral particles (VPs) of RhAd52 candidate vaccines following the indicated timeline. At week 12, mice were challenged with 10^4^ PFU of MA10 SARS-CoV-2 via the intranasal route. After challenge, a subset of mice was followed through day 4 post-challenge to monitor for signs of clinical disease. B) Relative body weight following MA10 SARS-CoV-2 challenge in mice immunized with the indicated vaccine regimens. Median value of each group is displayed. P<0.0001 indicates results of a one-way ANOVA analysis followed by Dunnett’s multiple comparisons, comparing vaccinated groups to the sham control group. C) Traces of relative body weight in individual mice, immunized with the indicated RhAd52 vaccine regimen, following challenge. For panels B-C, data are pooled from two similar experiments. N=5 mice/group for all regimens, with the exception of RhAd52.S Prime + Boost (N=10), RhAd52.S Prime Only (N=10), and Sham (N=13).

As expected, the sham control group exhibited significant weight loss following MA10 challenge, with a median loss of 15.2% of body weight at day 4 post-challenge (Fig 3B-3C). All vaccine regimens provided robust protection from clinical signs of infection in terms of body weights (P<0.0001, one-way ANOVA with Dunnett’s multiple comparisons test), with body weight generally remaining stable irrespective of the RhAd52 insert or whether a single or two-dose vaccine regimen was employed. Analyses of lungs revealed differences in the level of replication-competent virus detected in respiratory tract tissues among mice largely protected from weight loss (Fig 4A). In sham control mice at day 2 post-challenge, high levels of virus were recovered from lung, with a median titer of 3.1 x 10^7^ PFU/lung (Fig 4B). In contrast, two-dose regimens with full-length (i.e., S, S.PP) or truncated (i.e., S.dCT, S.dTM, S.dTM.PP) S immunogens provided a dramatic reduction in viral titer, with a greater than a 6 log drop in median titer. In nearly all mice in these groups, no replication competent virus was recovered from the lungs (i.e., PFU<100/lung). Immunization with two doses of the S fragment immunogens – RhAd52.S1 and RhAd52.RBD – restrained the level of virus in the lung, with a median titer of 3.9 x 10^5^ and 5.0 x 10^3^ PFU/lung, respectively. Similarly, in the single-shot groups, RhAd52.S and RhAd52.S.dCT provided significant but incomplete protection, reducing the viral burden in the lung to 3.3 x 10^2^ and 1.0 x 10^4^ PFU/lung, respectively. Finally, a single shot of RhAd52.S.PP dramatically reduced viral load, with no detectable viral outgrowth from lung tissues in 100% of mice. Of note, the S.PP insert previously proved optimal in nonhuman primates [24] and was advanced into clinical trials [19, 20].

**Fig 4.**
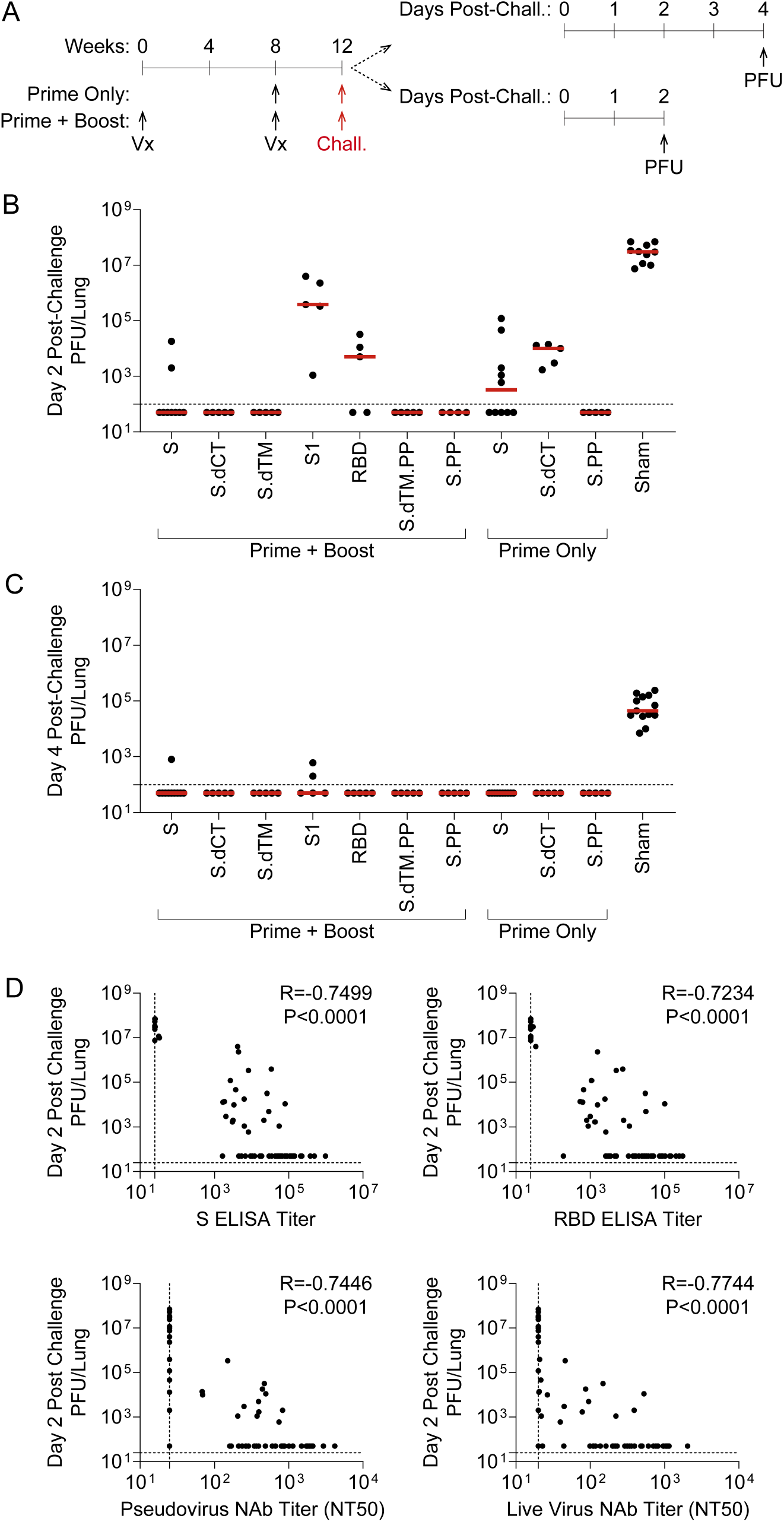
RhAd52 vaccine-elicited antibody responses link to restraint of viral replication in the lung following mouse-adapted SARS-CoV-2 challenge. A) Groups of mice were immunized with either a prime and a boost or a prime only of 10^9^ viral particles (VPs) of RhAd52 candidate vaccines. At week 12, mice were challenged with 10^4^ PFU of MA10 SARS-CoV-2 via the intranasal route. A subset of mice was monitored through day 4 post-challenge; at the terminal timepoint, lungs were harvested to measure virus via outgrowth assays to quantify plaque forming units (PFU) per tissue. The second subset of mice were followed through day 2 post-challenge for similar PFU assays at the time of peak viral replication. B-C) Quantification of PFU per lung at B) day two and C) day 4 post challenge. Median of each group indicated by the red line. N=4-5 mice/group for all regimens, with the exception of RhAd52.S Prime + Boost (N=9), RhAd52.S Prime Only (N=10), and Sham (N=10). D) Spearman correlation analyses of pre-challenge serum binding or neutralizing antibody responses with day two post-challenge viral titers in lung are displayed. For panels B-C, data are pooled from two similar experiments. In Panel E, data from all vaccine regimens are pooled to explore the relationship between pre-challenge binding and neutralizing antibody responses and virologic endpoint independent of the RhAd52 insert.

Similar analyses at day 4 revealed the level of virus in lung tissue was several logs lower (median 4.4 x 10^4^ PFU/lung) than at day 2 post-challenge (Fig 4C), consistent with our previous finding that tissue viral loads peak at day 1-2 post-challenge and gradually resolve over approximately 7 days [14]. Across vaccine regimens, virus levels were largely below the limit of detection of the outgrowth assay, with low levels observed in a subset of mice in the RhAd52.S and RhAd52.S1 two-dose groups. Together, these data suggest that all vaccines led to a reduction in respiratory tract tissue viral loads at the typical peak of infection as well as significantly decreased the persistence of virus in the lungs.

### Exploration of immune correlates of protection

We next evaluated possible correlations between vaccine-elicited immune responses prior to challenge and peak viral levels following challenge. A highly significant (P<0.0001) correlation was observed between pre-challenge S ELISA titers and peak (i.e., day 2) lung PFU (Fig 4D). Furthermore, the resulting Spearman correlation coefficient (R=-0.7499) suggests a strong inverse relationship between pre-challenge binding antibody levels and virologic outcome post-challenge. Similar highly significant (P<0.0001) inverse correlations were observed between three additional pre-challenge immunologic metrics and viral replication in the lung: i) RBD ELISA titers (R=-0.7234), ii) pseudovirus neutralization titers (R=-0.7446, and iii) live virus neutralization titers (R=-0.7744). Together, these data suggest that multiple vaccine-elicited humoral immune responses are inversely correlated with viral replication in respiratory tract tissue following MA10 SARS-CoV-2 challenge.

## Discussion

Our data indicate that RhAd52 vectors expressing SARS-CoV-2 S antigens elicit robust and protective humoral immune responses in mice. Based on baseline seroprevalence as well as the expanded global use of Ad5, Ad26, and ChAdOx1 vaccines [19, 20, 32–35], developing additional adenoviral vectors for COVID-19 vaccines is critical. This approach could be important for developing future boosting vectors or to tune the innate immune signatures induced [36]. Similar to our recent reports in hamsters [9], non-human primates [24, 28], and humans [19], a robust correlation was observed between binding and neutralizing antibody responses. Furthermore, although single-shot vaccines were highly protective, we observed increased immune responses using a homologous prime-boost strategy. In particular, the expansion of neutralizing antibody responses, as measured by both pseudovirus and live virus assays, in mice is encouraging, as this metric has emerged as a potential correlate of protection in hamster and non-human primate challenge models [9, 10, 21, 24, 28].

Importantly, the MA10 virus has previously been shown to drive significant clinical disease (i.e., weight loss) as well as replication localized in respiratory tract tissues, characteristics of interest for modeling severe COVID-19 disease. In contrast, nonhuman primate models for COVID-19 generally do not develop severe clinical disease [10-13]. The MA10 mouse model has proven useful for screening candidate therapeutics, but it remains relatively unexplored for testing vaccines [14-17]. The results from our challenge studies using RhAd52 vaccines suggest that candidate vaccines significantly protected against clinical disease and virus replication in lung tissue. However, only select immunization regimens drove full suppression of replicating virus in the lungs, as measured by viral outgrowth assays. Moreover, our data show that the recently-reported mouse-adapted virus MA10 exhibits robust humoral immune correlates of vaccine protection [9, 10, 21], which will prove useful in future studies of vaccines and other interventions using this model.

Future studies could further explore mechanistic correlates of protection, such as defining how vaccine candidates tune systemic pro-inflammatory cytokine secretion typically triggered by viral infection, as well as exploring the role of T cell responses in the context of MA10 challenge. No signs of disease enhancement (e.g., enhanced weight loss) were observed with sub-protective immune responses, an important finding due to concerns of antibody-dependent enhancement. Together, these data support the MA10 mouse-adapted virus as a tool to screen vaccine candidates. This approach could help to test novel immunogens, delivery systems, or dosing regimens, harnessing a relatively high throughput, tractable small animal model, wild-type mice. These studies could be employed to identify promising approaches to advance to large animal pre-clinical and, subsequently, early clinical trials. Moreover, similar mouse challenge models could be developed for the newly described SARS-CoV-2 variants of concern.

## Materials and Methods

### RhAd52 vectors

RhAd52 vectors were constructed with seven variants of the SARS-CoV-2 Spike (S) protein sequence (Wuhan/WIV04/2019; GenBank MN996528.1). Sequences were codon optimized and synthesized. Replication-incompetent, E1/E3-deleted RhAd52-vectors were produced in HEK 293B-55K.TetR cells as previously described [23], with the E1 region replaced by a transgene cassette encoding for the S sequence of interest. Vectors were sequenced and tested for expression before use.

### Animals and study design

Female BALB/c mice (The Jackson Laboratory) were randomly allocated to groups. Mice received RhAd52 vectors expressing different versions of the SARS-CoV-2 S protein or sham controls (*N* = 10 per group). Animals received a single immunization of 10^9^ viral particles (VPs) of RhAd52 vectors by the intramuscular route without adjuvant. In some cases, eight weeks later, mice received a homologous boost immunization. At indicated timepoints, peripheral blood was collected via the submandibular route to isolate serum for immunologic assays. For viral challenge, mice were administered 1 x 10^4^ PFU MA10 SARS-CoV-2 in a volume of 50μL via the intranasal route [14]. Following challenge, body weights were assessed daily. Subsets of animals were euthanized on days 2 and 4 post-challenge for viral outgrowth assays. All animal studies were conducted in compliance with all relevant local, state and federal regulations and were approved by the Beth Israel Deaconess Medical Center and University of North Carolina at Chapel Hill Institutional Animal Care and Use Committees.

### ELISA

S and RBD-specific binding antibodies were assessed by ELISA essentially as described [10, 28]. Briefly, plates were coated with 1 μg ml−1 of SARS-CoV-2 S protein (Sino Biological) or SARS-CoV-2 RBD protein (Aaron Schmidt, Massachusetts Consortium on Pathogen Readiness), diluted in 1× PBS, and incubated at 4 °C overnight. After incubation, plates were washed once with a wash buffer (0.05% TWEEN-20 in 1× PBS) and blocked with 350 μl of casein per well. The block solution was discarded after 2-3 hours of incubation at room temperature and plates were blotted dry. Three-fold serial dilutions of mouse serum in casein block were added to wells and plates were incubated for 1 hour at room temperature. Plates were then washed three times and rabbit anti-mouse IgG HRP (Jackson ImmunoResearch), diluted 1:1000 in casein block, was added to wells and incubated at room temperature in the dark. After 1 hour, plates were washed three times, and 100 μl of SeraCare KPL TMB SureBlue Start solution was added to each well. Development was halted with the addition of 100 μl of SeraCare KPL TMB Stop solution per well. The absorbance at 450 nm was recorded using a VersaMax microplate reader. ELISA endpoint titers were defined as the highest reciprocal serum dilution that yielded an absorbance > 0.2. The raw OD values were transferred into GraphPad Prism for analysis. A standard curve was interpolated using a sigmoidal four-parameter logistic (4PL) fit. To quantify the endpoint titer, the interpolation function was used to calculate the dilution at which the OD value would be equal to a value of 0.2.

### Pseudovirus neutralization assay

A SARS-CoV-2 pseudovirus expressing a luciferase reporter gene was generated in an approach similar to as described previously [10, 28, 29]. Briefly, the packaging construct psPAX2 (AIDS Resource and Reagent Program), luciferase reporter plasmid pLenti-CMV Puro-Luc (Addgene) and S protein expressing pcDNA3.1-SARS CoV-2 S.dCT were co-transfected into HEK293T cells using lipofectamine 2000 (Thermo Fisher Scientific). After 48 hours, supernatant was collected and pseudotype viruses were purified by filtration with a 0.45-μm filter. To determine the neutralization activity of the antisera from vaccinated animals, HEK293T-hACE2 target cells were seeded in 96-well tissue culture plates at a density of 1.75 × 10^4^ cells per well and incubated overnight. Three-fold serial dilutions of heat-inactivated serum were prepared and mixed with 50 μl of pseudovirus. The mixture was incubated at 37 °C for 1 hour before adding to HEK293T-hACE2 cells. 48 hours after infection, cells were lysed in Steady-Glo Luciferase (Promega) according to the manufacturer’s instructions. Neutralization titers were defined as the sample dilution at which a 50% reduction in relative light units was observed relative to the average of the virus control wells.

### Live virus neutralization assay

Live virus neutralization of sera was determined using a nanoLuciferase-expressing SARS-CoV-2 virus (SARS-CoV-2nLuc), bearing wild-type spike protein, as described [37, 38], with slight modification. Briefly, Vero E6 cells were seeded at 2 x 10^4^ cells per well in a 96-well plate 24 hours before the assay. 90 PFU of SARS-CoV-2-nLuc virus were mixed with serial diluted sera at 1:1 ratio and incubated at 37 °C for 1hour. An 8-point, 3-fold dilution curve was generated for each sample with starting concentration of 1:20. Virus and serum mix was added to cells and incubated at 37 °C + 5% CO2 for 48 hours. Luciferase activity was measured by Nano-Glo Luciferase Assay System (Promega) following manufacturer protocol using SpectraMax M3 luminometer (Molecular Device). Fifty percent neutralization titer (NT50) was calculated in GraphPad Prism by fitting the data points to a sigmoidal dose-response (variable slope) curve.

### PFU assay

Lung viral titers were determined by plaque assay. Briefly, right caudal lung lobes were homogenized in 1mL PBS using glass beads and serial dilutions of the clarified lung homogenates were added to a monolayer of Vero E6 cells and overlayed with a solution of 0.8% agarose and media. After three days, plaques were visualized via staining with Neutral Red dye and counted.

## Acknowledgments

We thank E. Hoffman, Z. Lin and R. Nityanandam for their generous advice, assistance, and reagents. We thank the veterinary and animal care teams at Beth Israel Deaconess Medical Center and University of North Carolina at Chapel Hill, particularly Andrew Sumski, Johanna Harvel, Melanie Harrington, Megan Brockett, Janet Veloz, and William Valle, for their assistance with animal care and transportation. We acknowledge support from the NIH (T32 AI007387) to L.H.T. We acknowledge support from Burroughs Wellcome Fund (Postdoctoral Enrichment Program Award), Howard Hughes Medical Insititute (Hanna H. Gray Fellowship), and NIH (T32 AI007151, F32 AI152296) to D.R.M. We acknowledge support from the NIH (AI145372) to L.E.G. This project was supported by the North Carolina Policy Collaboratory at the University of North Carolina at Chapel Hill with funding from the North Carolina Coronavirus Relief Fund (North Carolina General Assembly). We acknowledge support from the Chan Zuckerberg Initiative and the NIH (1U19 AI142759, 5R01 AI132178, AI100625, AI108197, HHSN272201700036I, and U54CA260543) to R.S.B. We acknowledge support from the NIH (CA260476, OD024917, AI149670, AI128751, AI129797, AI126603, AI124377), MassCPR, and Ragon Institute of MGH, MIT, and Harvard to D.H.B.

## Author Contributions

L.H.T., L.E.G., D.R.M., R.S.B., and D.H.B. designed and planned experiments. Z.F.L., J.L., and D.H.B. designed the RhAd52 vector vaccines. Z.F.L. and F.N. synthesized and characterized RhAd52 vectors. L.H.T. and S.H.M. performed the immunogenicity phase of the mouse studies. L.H.T., H.W., and K.A.M. performed ELISA assays. J.Y. and A.C. performed the pseudovirus neutralization assays. D.R.M. performed the live virus neutralization assays. D.R.M., L.E.G., and A.S. performed the MA10 challenge phase of the mouse studies. D.R.M., L.E.G., A.S., K.H.D., and S.R.L. performed the viral outgrowth plaque assays. L.H.T., L.E.G., and D.R.M. analyzed data and performed statistical analyses. L.H.T. and D.H.B. wrote the manuscript with input from and review by all authors.

